# Chromatin architecture and *cis*-regulatory landscape of the *DACT2-SMOC2* locus in the developing synovial joint

**DOI:** 10.1101/2022.10.06.511134

**Authors:** Karol Nowosad, Ewa Hordyjewska-Kowalczyk, Aneta Malesa, Adrian Odrzywolski, Rutger W. W. Brouwer, Petros Kolovos, Ilias Boltsis, Judith C. Birkhoff, Wilfred F. J. van IJcken, Frank G. Grosveld, Andrea Conidi, Danny Huylebroeck, Przemko Tylzanowski

## Abstract

**Background:** Synovial joints form in several steps, starting with the formation of an interzone, a condensation of mesenchymal cells at the sites of prospective joints. Despite the identification of multiple factors essential for formation of interzone, little is known about the regulation of their spatio-temporal gene expression during that process in limb development. Here, we investigated the *cis-* regulatory landscape of the Wnt-modulator encoding genes *DACT2* and *SMOC2*, both expressed in the forming joint interzone.

**Results:** Mechanically collected interzone and phalange samples, respectively, from chick embryos were found to express acknowledged marker genes (*GDF5* and *MATN1*), as well as *DACT2* and *SMOC2*. Using Targeted Chromatin Capture (T2C) we characterized the 3D chromatin structure of a ~3.45 Mb-long region encompassing *DACT2* and *SMOC2*, which revealed differences at sub-TAD level between interzones and phalange. We identified candidate enhancers (CEs) based on H3-histone marks (H3K427ac and H3K4me1) located in close proximity to the promoters of *DACT2* and *SMOC2*, and further documented these CEs in a zebrafish enhancer assay.

**Conclusions:** Our approach yields new insight into the regulation, in dynamic chromatin context, of two Wnt-signaling modulatory genes during synovial joint induction.

## Introduction

The developing vertebrate limb is a frequently used model system to study the genetic and molecular control mechanisms of tissue/organ induction and subsequent patterning. Limb skeletal elements are derived from lateral plate mesoderm, while the limb muscle components originate from cells of the dermomyotome, a part of the transient, segmented somite formed in the paraxial mesoderm. During limb outgrowth, the mesenchymal cells undergo condensation followed by chondrogenic differentiation, resulting in the formation of a transient cartilage scaffold. The concomitant tissue patterning along the three limb axes determines the location and shape of the future bones^1–3^. At the onset of chondrogenic differentiation, these cells express *COL2A1* (α(II)-collagen) and *MATN1* (Matrilin-1)^4^, followed by *COL10A1* (α(X)-collagen), which is specific for hypertrophic chondrocytes^5^. Eventually, this hypertrophic cartilage becomes vascularized and is replaced by bone^6^. In contrast, the cells from the interzone region, located between the ends of future skeletal elements, are involved in the development of synovial joint structures, which include articular cartilage, menisci, ligaments and the synovium itself^7–10^. Interzone cells express acknowledged marker genes such as *GDF5* (Growth and Differentiation Factor-5), *ATX/ENPP2* (Autotaxin), *WNT9B* (Wnt Family Member 9B) and *ERG* (an ETS family transcription factor)^9,11,12^.

Significant efforts have been made to analyze the transcriptome of developing synovial joints^13–16^, but enhancer-driven regulation of gene expression during joint formation remains underexplored. Our laboratory studies the induction of the synovial joint. As part of these efforts, we study the role of a secreted Wnt/BMP signaling modulator, *SMOC2*, isolated from articular cartilage^17–20^. *SMOC2* mRNA is expressed in several tissues, including the developing synovial joint of the E14.5 mouse embryo^21^. *DACT2* (Dishevelled binding Antagonist of beta Catenin 2), one of the *SMOC2* neighboring genes on the same chromosome, is also expressed in the interzone region^22^. The co-localization of these two genes in the genome is conserved among human, mouse, chicken, and zebrafish with species-specific differences of the length of the intergenic region (from 75 to 150 kb). *DACT2* negatively regulates Wnt-β-catenin signaling by disrupting the β-catenin:LEF1 complex in the nucleus^23^. It also modulates YAP/TAZ signaling by preventing nuclear accumulation of Yes-Associated Protein (YAP)^24^, a transcription factor (TF) involved in the negative regulation of the gene encoding the BMP-subgroup ligand *GDF5*^25^. Since both *SMOC2* and *DACT2* are co-expressed during joint formation, and are genomically separated in a “head-to-head” configuration by an intergenic region, we hypothesized that they may share transcriptional regulatory elements.

Distant genomic enhancers may control target gene expression in spatiotemporal manner, orchestrating cell-type specific pattern of genes^26^. Enhancers co-regulate RNAPol2-based gene transcription by bridging TFs and relevant co-factors with the promoter-proximal region of their target gene(s), likely via DNA-looping^27^. Importantly, the organization of chromatin architecture into topologically associating domains (TADs), delineated by borders enriched for DNA-binding sites of the multiple zinc-finger protein CTCF, was shown to promote intra-TAD enhancer-promoter contacts and insulate inter-TAD interactions^28–31^. Therefore, enhancers function mainly in *cis* and within TADs. Such enhancer activity also correlates with overall chromatin accessibility, established in part by nucleosome positioning and dynamics, and is secured by histone modifications (e.g., H3 acetylation (ac) and methylation (me), such as H3K27ac and H3K4me1 signatures)^32^.

Chromosome conformation capture (3C) and its derivatives (4C, 5C, Hi-C) are powerful tools for investigating enhancer-promoter interactions in higher-order 3D chromatin context^33–36^. These techniques however provide low to moderate resolution only, or require large or deep sequencing efforts. A recently developed technique, i.e. Targeted Chromatin Capture (T2C), a 3C-based variant, addresses these limitations and permits to obtain high-resolution data at affordable sequencing cost^37–40^. Another advantage of T2C is the targeted enrichment of short genomic regions of interest by hybridization with custom oligonucleotide probes, which increases signal-to-noise ratio.

Recently, we developed an atlas of candidate enhancers (CEs) active in interzone and adjacent phalange, and associated these CEs with genes upregulated in the interzone^41^. We also linked such CEs to genes known as crucial in synovial joint hypermobility and osteoarthritis, as well as phalange malformations^41^. This CE atlas is serving as resource for identifying and validating enhancer-controlled synovial joint and phalange formation, but also provided us a new starting point to study the regulation of *DACT2* and *SMOC2* expression at the onset of joint formation. Using a combination of integrative analysis of chromatin 3D architecture, CEs defined by enrichment of histone modifications (H3K27ac and H3K4me1), and an enhancer assay in zebrafish larvae, we report here the identification of multiple CEs associated with *DACT2*, one of which is shared with *SMOC2*, in the developing joint.

## Results

### Micro-dissection of interphalangeal joint interzones

To collect the samples for T2C, we dissected and separated joint interphalangeal interzones and the adjacent proximal part of phalange from digit-3 of the chick embryo hindlimb at HH32 (see Materials and Methods). Validation of the purity of the dissected tissues was done by RT-qPCR analysis of acknowledged marker mRNAs for interzone (*GDF5*) and phalange (*MATN1*), and showed that both samples were successfully dissected and separated (**Figure 1A**). Previously, we carried out RNA-Sequencing (RNA-Seq) on such samples and showed significant upregulation of steady-state *DACT2* mRNA in dissected interzones as compared to phalanges, whereas *SMOC2* mRNA displayed high variation between biological replicates of interzone samples. However, a trend towards upregulation in interzone tissue was observed ^41^.

**Figure 1.**
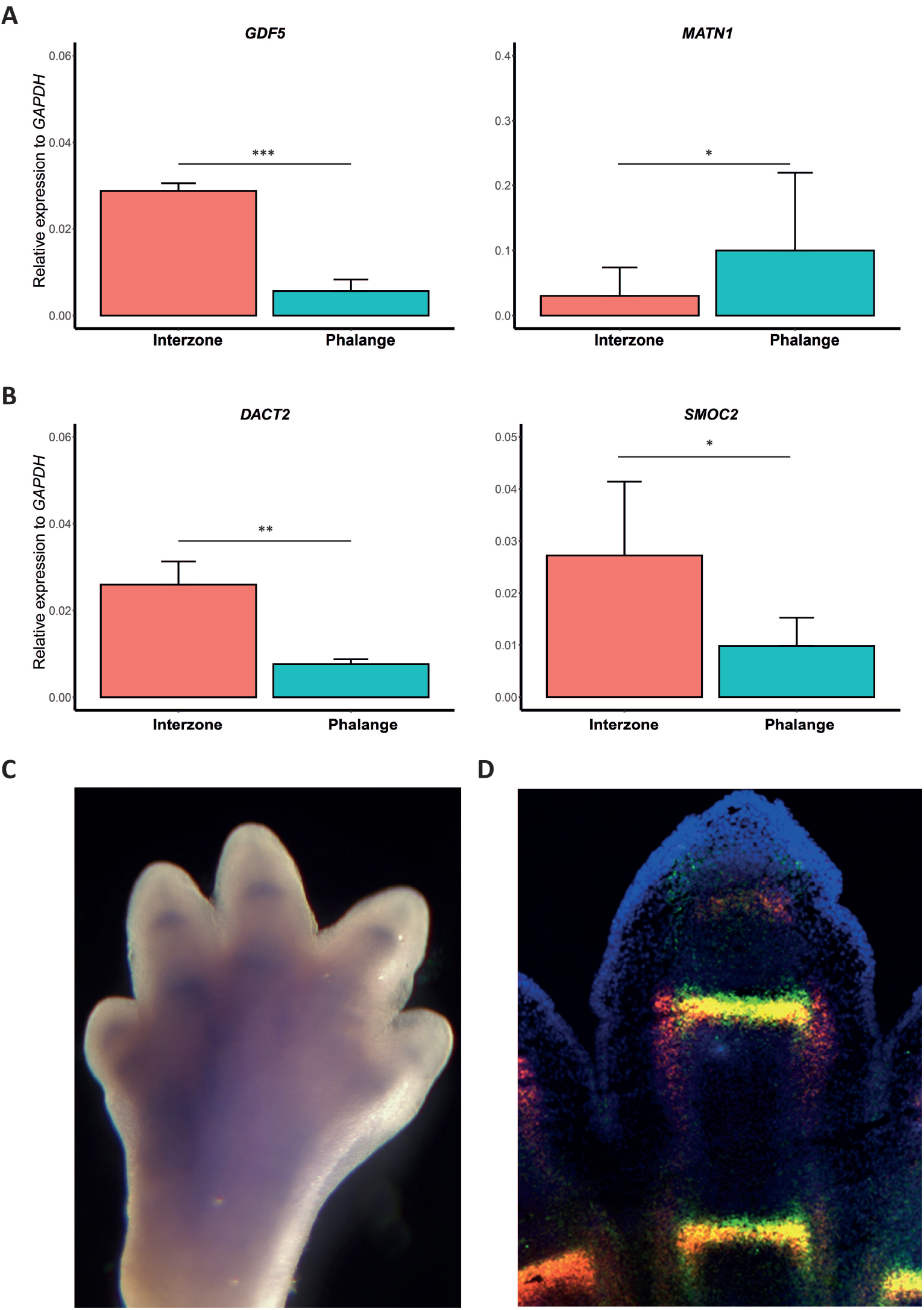
Expression of *DACT2, SMOC2 and* interzone (*GDF5*) and phalange (*MATN1*) marker genes. (A) The *GDF5,MATN1*, and (B) *DACT2, SMOC2* steady-state mRNA levels, determined by RT-qPCR, after dissection of phalanges and interzones from chick embryo hindlimb digit-3 at stage HH32 (see Experimental procedures). All data was normalized to expression of *GAPDH*. The p-value is based on T-test. (C) Whole-mount *in situ* hybridization on the autopod of a E14.5 mouse forelimb. The blue staining depicts *Smoc2* mRNA expression. (D) Dual *in situ* hybridization in a digit of a E14.5 mouse forelimb. In red: *Gdf5* mRNA, green: *Smoc2* mRNA, yellow: overlap between *Gdf5* and *Smoc2* expression, blue: nuclear counterstain using DAPI.

RT-qPCR on the dissected tissues used in the present study showed significantly higher expression of both *DACT2* and *SMOC2* in dissected interzones as compared to phalanges (**Figure 1B**). To document the *Smoc2* expression domain, we additionally performed whole-mount (**Figure 1C**) and dual *in situ* hybridization (using also *Gdf5* anti-sense probe; **Figure 1D**) in forelimb digits from E14.5 mouse embryos. The *Smoc2* expression was found to increase in the interzone, and partially overlapped with joint-specific *Gdf5* transcripts, but extended distally, beyond the *Gdf5* mRNA expression domain. In summary, *Smoc2* mRNA is expressed in the developing synovial joint.

### Chromatin organization of the *DACT2 – SMOC2* genomic region during joint formation

Following the tissue isolation, the interzone and phalange 3D chromatin structure within a ~3.45 Mb region (chicken chr3: 40,15-43,6 Mb) encompassing *DACT2* and *SMOC2* was determined by low-T2C, using a protocol optimized for lower cell numbers^39^. First, we validated the efficiency of targeted fragment enrichment within the region of interest by quantification of paired reads mapped to both the whole genome and the target region. This showed that such reads for this target region were enriched in both tissues (**Table S1)**. Next, we removed the self-ligated/non-digested regions and calculated the *cis/trans* interactions (**Figure 2A**) and fragment density distribution together with median resolution (**Figure 2B**). The analysis of *cis* and *trans* interactions revealed that the T2C data contained much more *cis* interactions, further suggesting that these data are of good quality. The comparison of fragment length distributions showed that these did not change with tissue, while the median resolution (~570 bp) revealed that both interzone and phalange T2C datasets were generated at sub-kb resolution.

**Figure 2.**
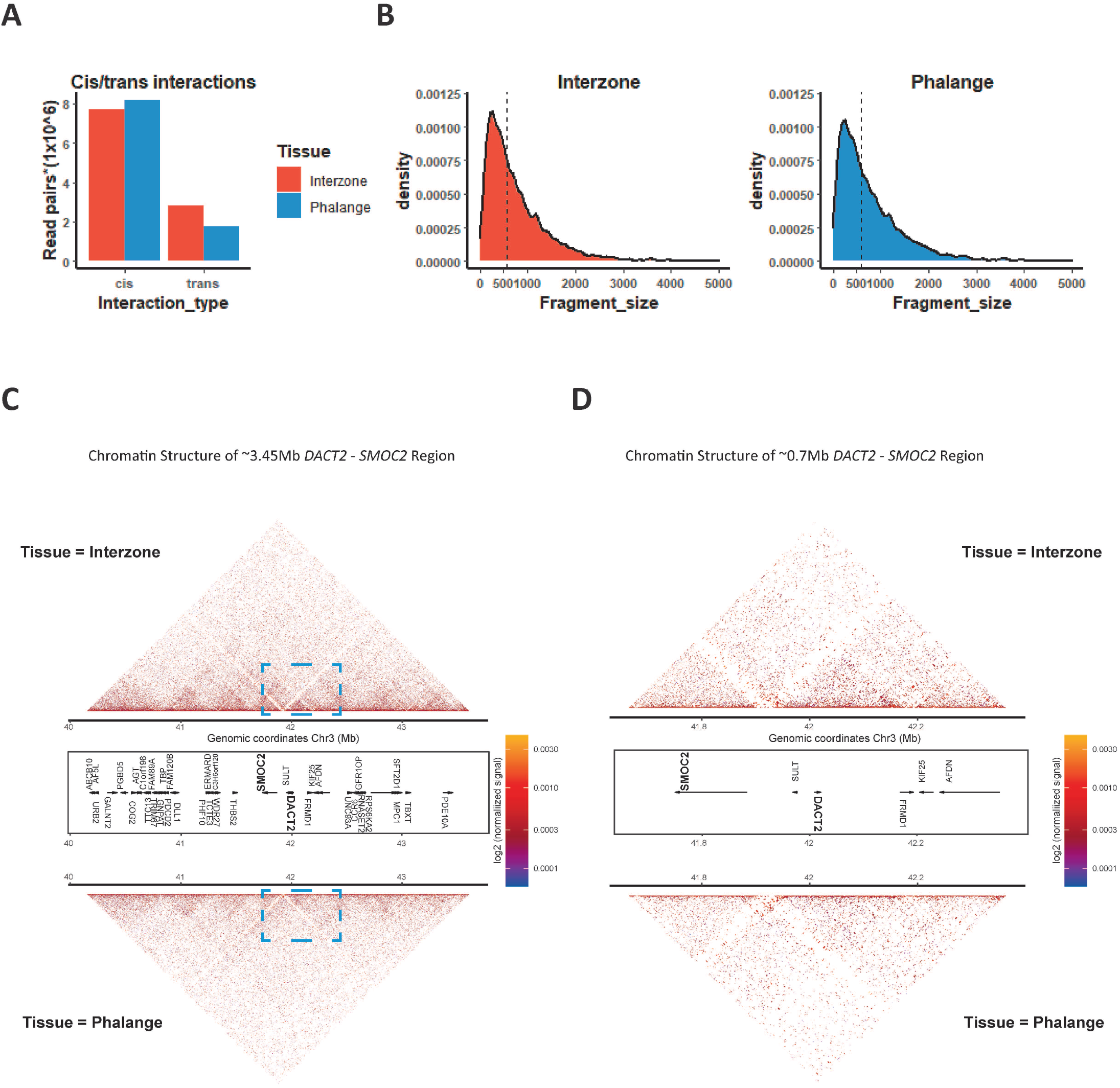
Characterization of the 3D chromatin structure of *DACT2-SMOC2* genomic region as determined by T2C. (A) Number of *cis* and *trans* targeted read-pairs in the proximity matrix. (B) Density plot of fragment distribution within *DACT2–SMOC2* ~3.45 Mb genomic region. Dashed line presents median fragment size. (C) T2C-interaction map (restriction fragment resolution) for the ~3.45 Mb-long *DACT2-SMOC2* genomic region (chicken chr3: 40,15-43,6 Mb) for separated interzone tissue (upper triangles in panels C and D) and phalange (lower triangles). The dashed rectangle marks the zoomed region shown in panel D. (D) The zoom-in genomic region chr3: 41,6-42,3 (~0.7 Mb-long) presents the chromatin architecture at inter-TAD level. Tissues are marked as in panel C.

The resulting T2C interaction maps for the chicken *DACT2-SMOC2* region (**Figure 2C**) confirmed the global organization of the chicken genome into TADs, in line with a recent Hi-C study^42^. The maps also confirmed the hypothesis that TADs are conserved among tissues, in this case between interzone and phalange. The quantification of unique fragment pair proximities within the region of interest showed that the interzone dataset contains a higher number of fragment pairs when compared to the phalange dataset (**Table S1**). This was reflected in more pronounced interzone-specific patterns of fragment proximities in the T2C interaction maps compared to those in phalange, which presented with a relatively more diffused pattern of such proximities (**Figure 2CD**, with the entire *DACT2-SMOC2* 3.45 Mb-long region in **panel C**, and a zoomed-in map for a 0.7 Mb segment in **panel D**). To characterize the intra-TAD chromatin organization, we analyzed the chicken chr3:41,6-42,3 Mb zoomed-in region (~0.7 Mb), showing differences between interzone and phalange within this region (**Figure 2D**). The divergent pattern of the intra-TAD interaction between the two dissected tissues is most likely caused by DNA-looping of tissue-specific regulatory elements to their target gene/s, for instance *DACT2* and *SMOC2*, which is our focus for further studies here.

We extracted genomic regions located in spatial proximity to the gene promoters (defined as a 5 kb-long segment, −2.5/+2.5 kb flanking the transcriptional start site, TSS), and averaged the signal from T2C to the length of the promoter region by binning the data, using a bin size of 5 kb (**Table S2**). Subsequently, we quantified the number of *cis*-proximities and showed that the promoter of *DACT2* and *SMOC2* presented higher percentages of tissue-specific interactions as compared to common interactions, and this in both dissected tissues (**Figure 3**). Interestingly, for both genes the ratio of tissue-specific vs. consensus *cis*-proximities was higher for the interzone than for the phalange (**Figure 3**).

**Figure 3.**
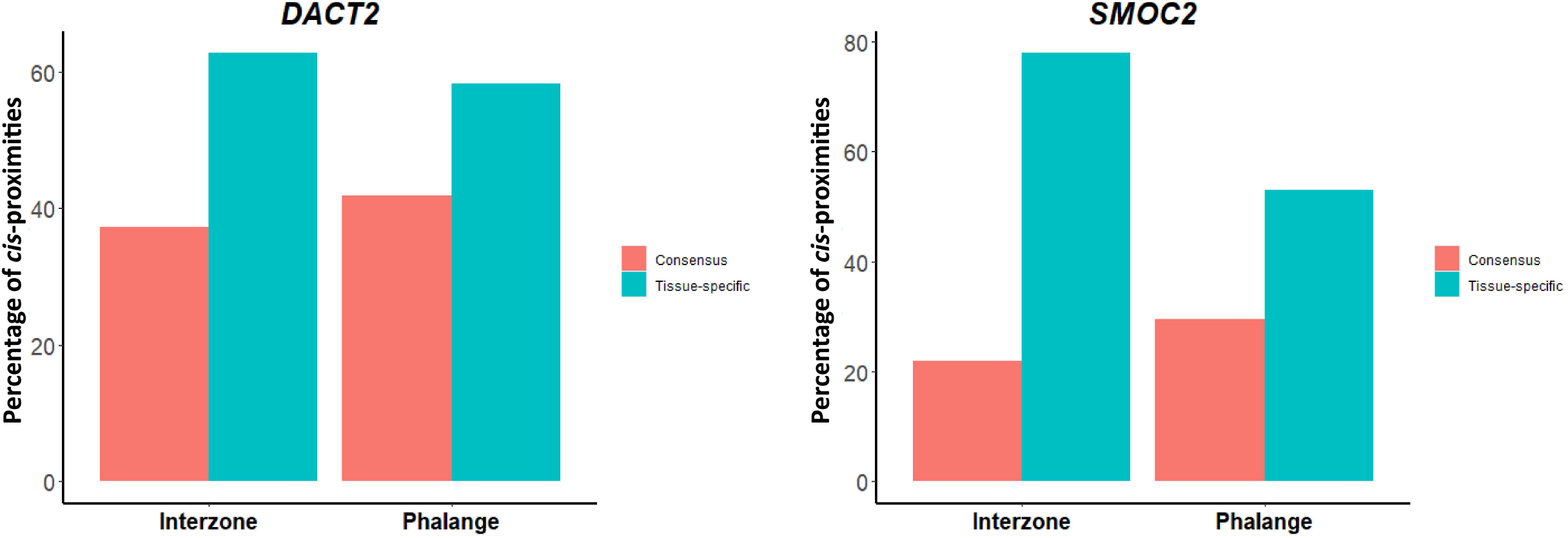
Quantification of *cis*-proximities of *DACT2* and *SMOC2* promoters. The genomic regions located in spatial proximity to the gene promoters (−2.5/+2.5 kb from TSS) were quantified for *DACT2* (left) and *SMOC2* (right). Further, cis-proximities have been divided into separated groups (tissue specific, consensus for both tissues) and presented as a percentage of all interactions identified per tissue.

### Identification of *DACT2* and *SMOC2* candidate enhancers

Next, to identify CEs in the ~3.45 Mb-long *DACT2-SMOC2* region, we screened our CEs atlas^41^ for these enhancers that are active in interzone and phalange. This analysis showed either low or no enrichment of *cis*-proximities between CEs and promoters of *DACT2* and *SMOC2*. However, this could be explained by the fact that our enhancer atlas contains only CEs that are conserved between chicken, human and mouse. Therefore, we had to apply a less stringent species conservation context for the analysis of CEs in the study here. We subsequently reanalyzed the original species-specific ChIP-Seq profiles of H3K27ac and H3K4me1 that were used to define the CEs in our atlas^41^. Using this strategy for the ~3.45 Mb-long *DACT2-SMOC2* region, we identified 53 interzone CEs and 35 phalange CEs, defined here as regions enriched for both H3 marks (**Table S3**). To further characterize chicken CEs, we carried out differential analysis of H3K27ac marks to investigate which of these CEs are located in differentially acetylated regions (DacRs). This revealed that the ~3.45 Mb-long *DACT2-SMOC2* region in chicken contains 59 DacRs, 45 of which were highly enriched in interzone as compared to phalange (**Table S4**). H3K27ac is a mark for active enhancers, and the majority of DacRs were enriched in interzone, suggesting that the *DACT2-SMOC2* region contains more differentially active enhancers in the interzone than phalange region. Importantly, intersection of CEs with DacRs identified that 33 CEs in interzone and 19 CEs in phalange, respectively, were located in DacRs (**Table S5**).

To identify CEs that regulate *DACT2*, we screened all interacting regions with *DACT2* promoter and intersected them from the aforementioned 88 CEs (53 interzone + 35 phalange). This permitted to select 7 CEs with a high T2C-score and enrichment for both H3 marks as well (**Figure 4; Table S6**). Three of these (named CE1, CE3, CE5) were mapped in proximity to *DACT2* exclusively in interzone sample (**Figure 4AB**), whereas the remaining other 4 (named CE2, CE4, CE6, CE7) were found close in 3D space to the *DACT2* promoter in both tissues (**Figure 4B**). Intersecting these selected CEs with DacRs revealed that CE1-4 and CE7 were located in DacRs, whereas CE5-6 did not present with significant differences in this acetylation level (**Figure 5A**).

**Figure 4.**
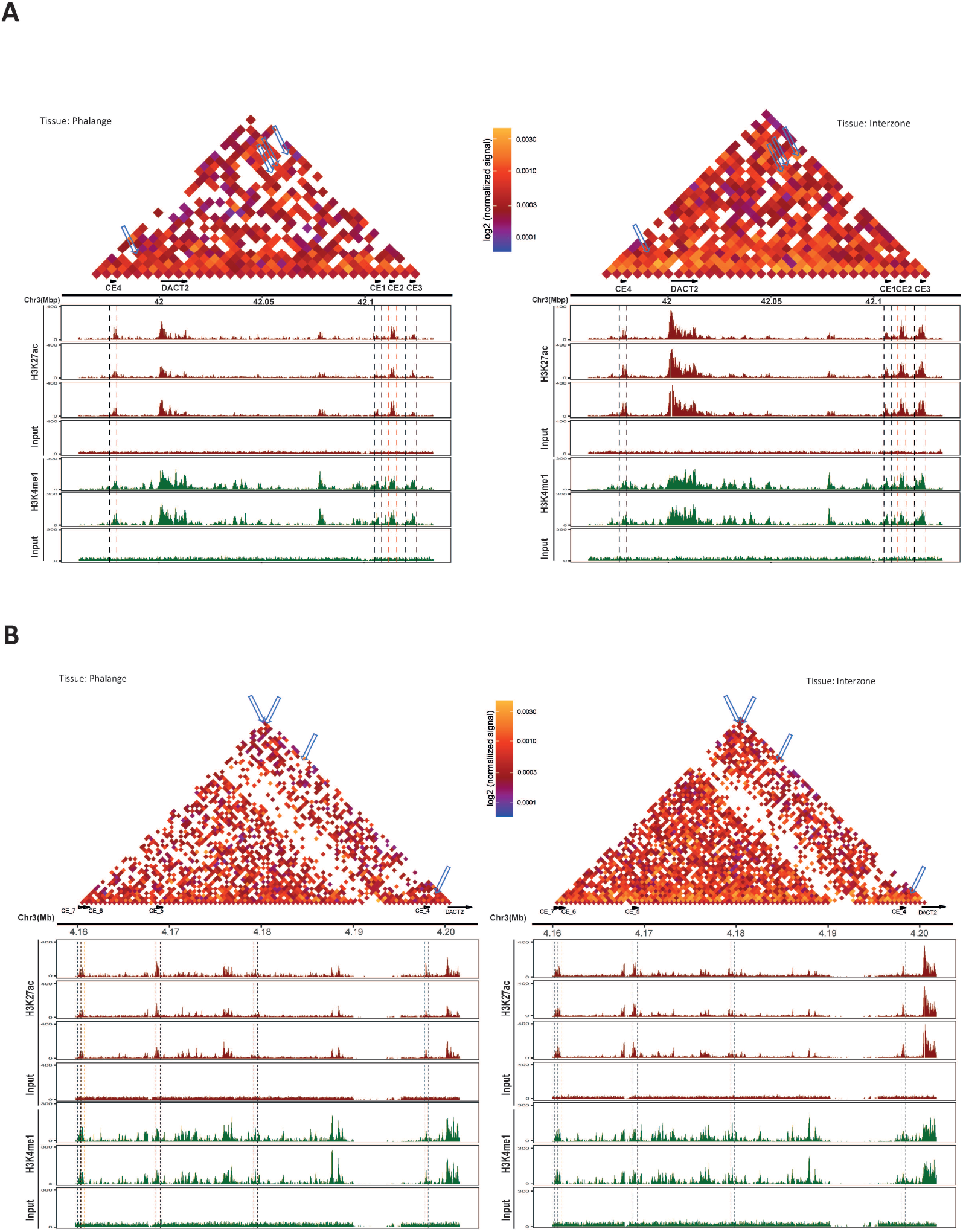
Identification of *DACT2* CEs. (A) T2C interaction maps (bin size = 5 kb) encompassing identified CE1, CE2, CE3 and CE4 and *DACT2. Cis*-proximities between enhancers and *DACT2* promoter are indicated by blue arrowheads. Dashed lines show selected CEs regions enriched in histone modification. The H3K27ac tracks are highlighted in dark red. The H3K4me1 tracks are marked by dark green. ChIP-Seq inputs are marked by black. (B) T2C interaction maps (bin size = 5 kb) presenting proximities (pointed by blue arrows) between *DACT2* and CE4-CE7. CEs and histone modification tracks are marked as in panel A.

**Figure 5.**
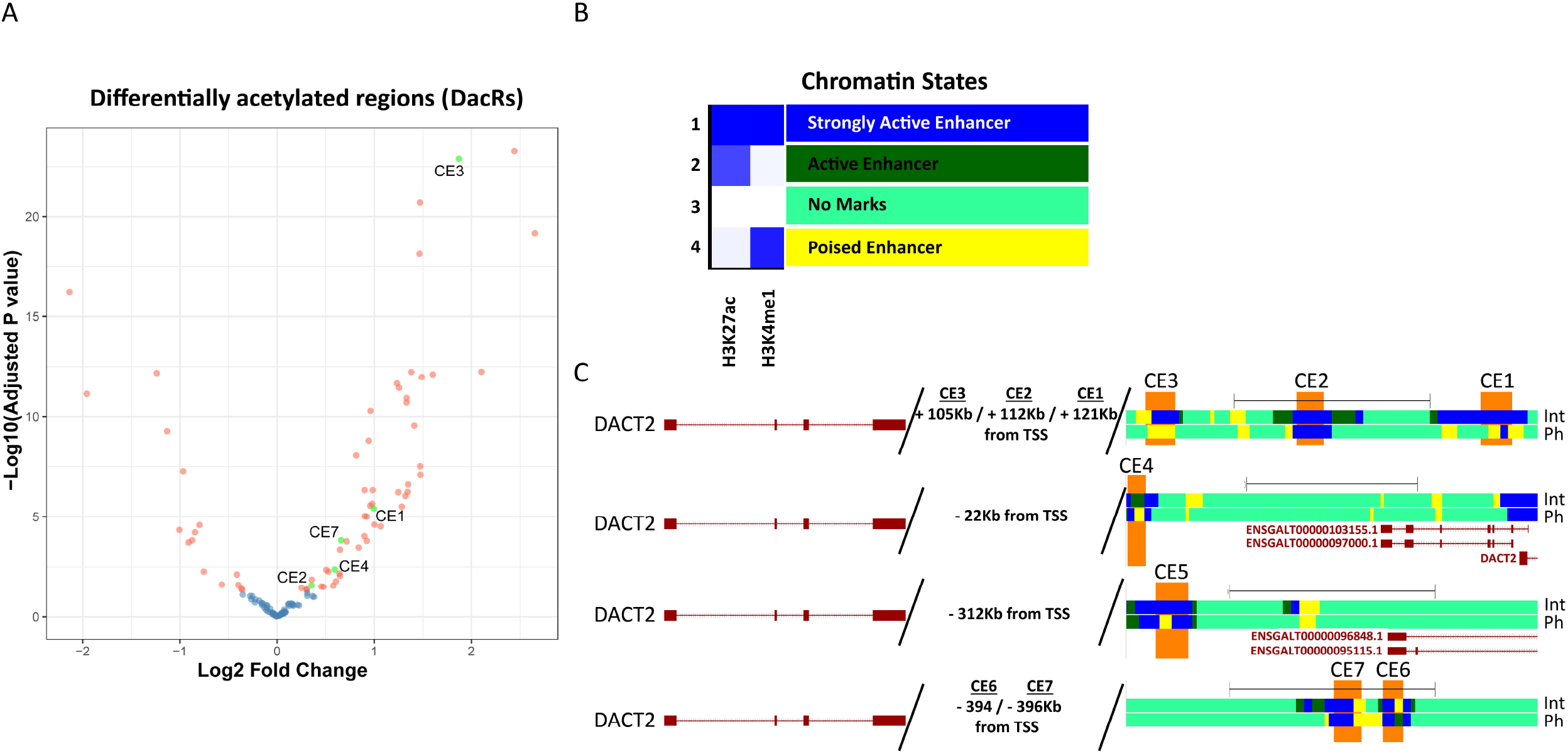
Further characterization of *DACT2* CEs. (A) Plot presenting differentially acetylated regions (DacRs). The plot was generated using DiffBind based on the enrichment of H3K27ac in interzone and phalange samples. Blue dots represent DAR with FDR > 0.05, salmon-color dots mark significantly differentiated DAR (FDR <0.05). Green dots depict DacRs encompassing selected CEs. (B) A 4-chromatin state model obtained using interzone and phalange histone ChIP-Seq data. The chromatin state model was generated using ChromHMM based on all sample replicates, including input controls. The states were annotated using the model emission probabilities (**Table S7**) visualized by the intensity of blue color within the heatmap. (C) The UCSC genome browser view of interzone and phalange states around of selected CEs. Colors of chromatin states correspond to state annotation in panel B.

Next, we analyzed whether the seven selected CEs change enhancer state from strongly active (i.e. enriched in H3K27ac and H3K4me1) to poised enhancer (enriched in H3K4me1 only). For this, we estimated the chromatin state signatures with ChromHMM tool, which uses a multivariate hidden Markov model (HMM) based on ChIP-Seq data from both H3 marks. Four such chromatin state signatures have been identified as signature A (strongly active enhancer), B (active enhancer), C (poised enhancer) and D (no enhancer marks), respectively (**Figure 5B**; **Table S7**).

This comparison revealed that the major part of the CE1 region changed from strong to poised enhancer/no enhancer state, like the entire CE3 region did. In contrast, CE2 (located between CE1 and CE3) was identified as strongly active enhancer in interzone (Int) and phalange (Ph in **Figure 5C**, top line). Further, CE4 switched from active to poised enhancer in the majority of its region, and CE5 from strongly active to poised enhancer only in a part of its region (**Figure 5C**, 2nd and 3rd line from the top). A part of CE6 switched from poised enhancer in interzone to active enhancer in phalange; in contrast, CE7 presented with the same chromatin state in both tissues (**Figure 5C**, bottom line). These data support the hypothesis that the chromatin state between interzone and phalange tissue during joint formation in the embryo is different and dynamic around the *DACT2* locus.

Similar investigation of CEs possibly involved in the regulation of the *SMOC2* locus revealed that CE7 is located in spatial proximity to the *SMOC2* promoter, suggesting that CE7 regulates both *SMOC2* and *DACT2*. Further analysis of our T2C data indeed revealed that both genes are brought in spatial proximity to CE7 (**Table S2**). For visualizing the *cis*-proximities between each gene and the putative common enhancer CE7, we generated virtual 4C tracks based on our T2C data, and combined this with H3K27ac and H3K4me1 enrichment analysis. This operation, using virtual 4C *DACT2/SMOC2* promoters as viewpoints, confirmed the presence of *cis*-proximities between the promoter segment of both genes and CE7. Also, this 4C approach showed that both gene promoters are located in spatial proximity (**Figure 6**).

**Figure 6.**
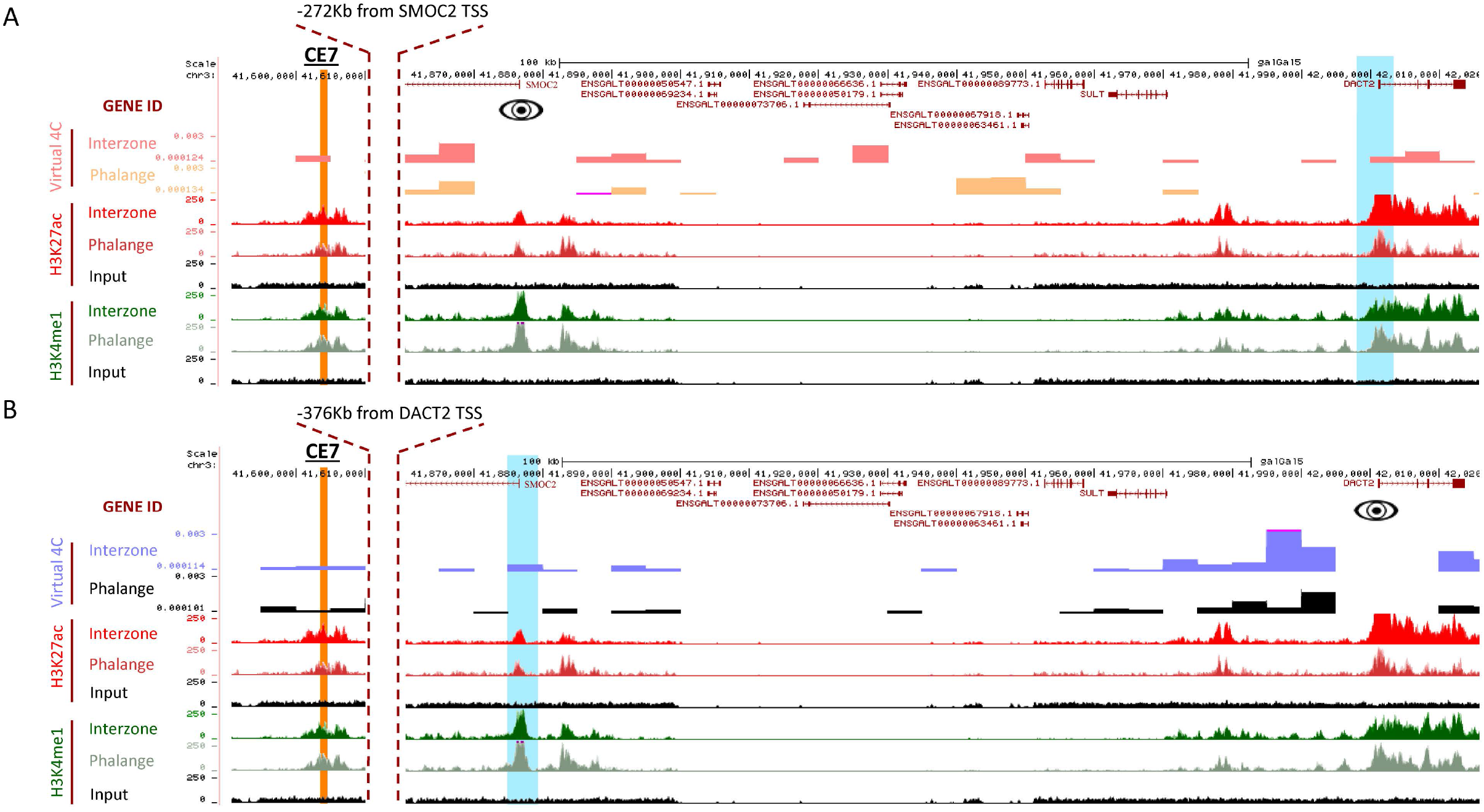
Characterization of virtual 4C for *DACT2* and *SMOC2*. (A) The virtual 4C was generated using the T2C data for interzone and phalange samples. The genomic region coordinates are indicated in the top lane, the height of peaks corresponds to signal value. The signal value was averaged using bin size = 5 kb. The virtual 4C for *SMOC2* as a viewpoint (“eye” symbol) revealed the loop between *SMOC2* and CE7 (dark orange), and that and *SMOC2* and *DACT2* promoter (highlighted by blue) are in spatial proximity. The Virtual 4C track for interzone marked by salmon-color and phalange by orange. The H3K27ac enrichment tracks are marked by red, and H3K4me1 by green. Input control tracks are marked by black. (B) Characterization of virtual 4C tracks (blue for interzone and black for phalange) generated for *DACT2* as a viewpoint (“eye” symbol) showed cis-proximity between *DACT2* and CE7 were (highlighted by dark orange). Also, in this case the *DACT2 and SMOC2* promoter (marked in blue) have been identified as located in spatial proximity. The H3 histone marks tracks colored as in panel A.

### *In vivo* testing of candidate enhancers using a zebrafish enhancer assay

To test *in vivo* activity of selected *DACT2* CEs we used a zebrafish enhancer assay using a Zebrafish Enhancer Detection (ZED) vector containing two expression cassettes^43^. One of the cassettes contains a Green Fluorescent Protein (GFP) cDNA driven by the minimal *GATA2A* promoter, which can be activated by an enhancer placed upstream of it. A second cassette serves as an integration control where Red fluorescent protein (DsRed) mRNA is placed under the control of Cardiac Actin Promoter, which is active in the zebrafish heart and somites. Embryos were injected at the 1-cell stage and monitored under a fluorescent microscope every 24 hours (*data not shown*). Most prominent production of GFP for each of our selected 7 CEs was observed 96 hours post fertilization (hpf). The specific GFP signals presented in at least 30% of embryos analyzed were considered specific.

We found the most prominent activity of the selected CEs in the regions of forebrain, midbrain, otolith and jaw cartilages (**Figure 7A**), where *DACT2* is also expressed in zebrafish^44^. We noticed that CE1 and CE2 were active in the branchial arches of the developing larvae, partially sharing regulatory information, although with varying fluorescence intensities (**Figure 7D**). Further, the low intensity GFP signal for CE3 was observed around the otic vesicle (**Figure 7D**). CE6 was active in the forebrain and around the otic vesicle (**Figure 7E**), however its signal in the forebrain region sporadically overlapped with ZED auto-fluorescence. So, the regulatory role of CEs in this specific area cannot be fully determined, although the detected CE signal was invariably stronger than in the group injected with empty (without CE) vector (**Figure 7C**). Contrary, we could never detect signals from CE4, CE5 and CE7 (**Figure 7E**). Importantly, the localization of GFP for CEs1-3 and CE6 in developing larvae resembled the pattern of RNA *in situ* hybridization of *DACT2* in zebrafish larvae^44^. Overall, these results show overlap between the *DACT2* expression and its CEs when tested *in vivo*, suggesting the involvement of the analyzed CEs in the formation of those structures.

**Figure 7.**
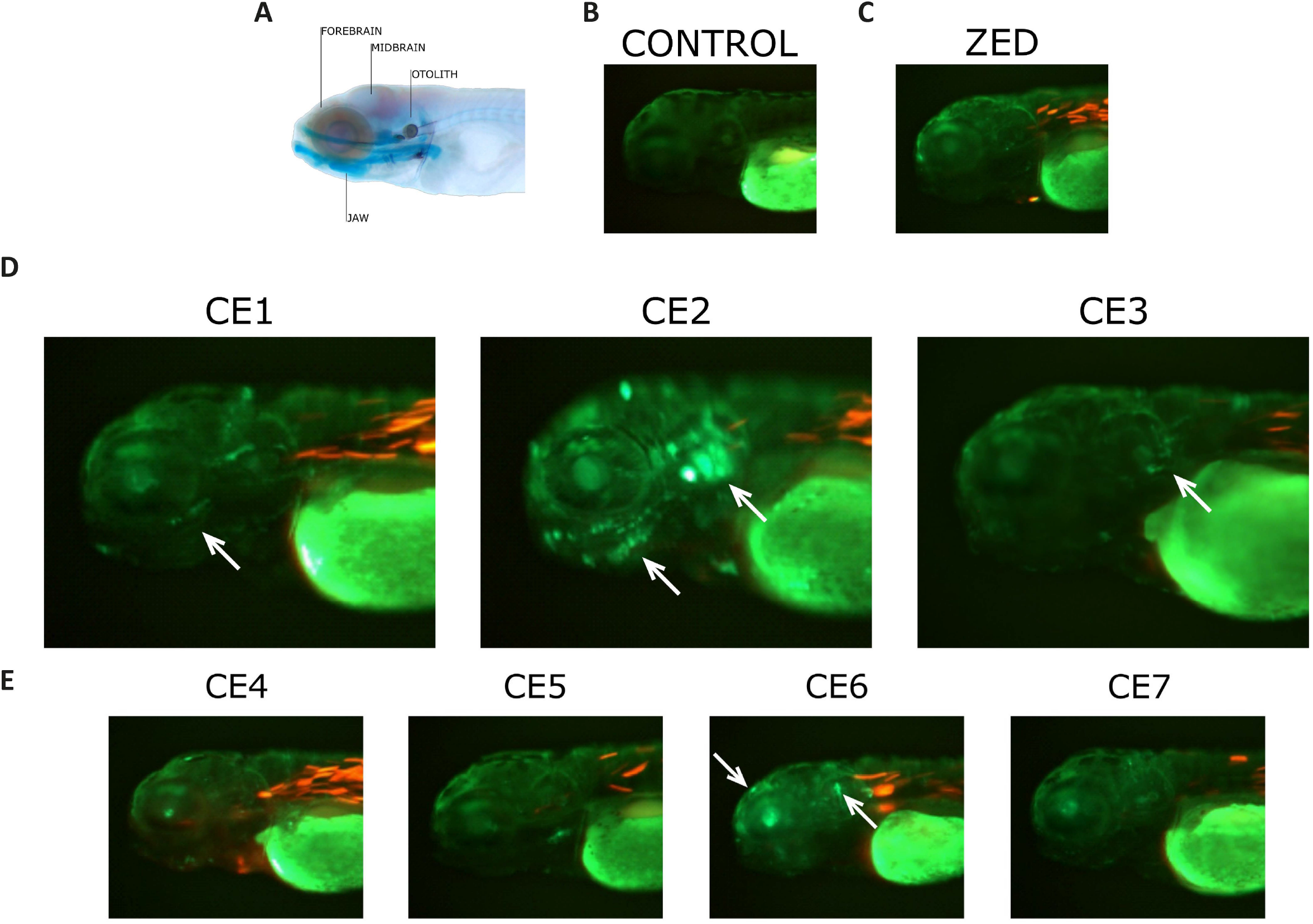
CEs activity in developing zebrafish larvae. (A) Zebrafish larvae at 120 hpf stained with Alcian Blue (cartilage) and Alizarin Red (bone) to reveal the skeletal structures. (B) Uninjected zebrafish larvae at 96 hpf observed in the green channel, revealing the autofluorescence. The autofluorescence in the red channel was not detected (data not shown). (C) Empty vector injected larvae at 96 hpf. (D) Representative merged fluorescent images showing the activity of enhancer CE1, CE2 or CE3-driven GFP and DsRed integration control. The arrow in CE1 points to the developing jaw, in CE2 to the developing jaw and otic vesicle and in CE3 to the otic vesicle only. (E) Representative images showing activity of tested CE4-7 (GFP), DsRed indicates integration control. The arrow in CE6 points to the otic vesicle and forebrain.

## Discussion

Limb development, including patterning and cell differentiation is subject to precise and dynamic spatiotemporal transcriptional control. There are many examples describing the intricate regulatory circuits for limb-relevant genes such as *SHH, HOXD* cluster or *GDF5*^45–48^. The deregulation of such genes, caused for instance by mutation in enhancers or by chromosomal rearrangements, leads to limb deformities^49–52^. In this study, we investigated the role of chromatin architecture and enhancers in the expression of two genes co-expressed during joint development, *DACT2* and *SMOC2*. We opted for the physical dissection of interzone tissue instead of one based on the isolation of *GDF5*-positive (+) cells, because not all cells during early stages of joint formation are convincingly *GDF5*+^16^. Dissected interzones presented significantly higher expression of *GDF5*, *DACT2* and *SMOC2* as compared to adjacent phalange, consistent with publicly available^21,22,48^ as well as our own *in situ* hybridization studies.

To explore the potential transcriptional co-regulation of both genes, we analyzed the chicken chr3:40,15-43,6 genomic region encompassing the *DACT2-SMOC2* loci, focusing on the chromatin architectural dynamics for joint interzone and the adjacent phalange. Based on T2C experiments, we detected DNA-loops within the TADs, which were predominantly tissue-specific. This is in agreement with Hi-C data showing that TADs tend to be conserved between tissues and cell types, in contrast to intra-TAD interactions, which within different cell types can be dynamic^29,42^.

Multiple studies point out the role of enhancers in co-shaping chromatin architecture^29,53,54^. Using our recent data from ChIP-Seq for H3K27ac and H3K4me1^41^ we identified 88 CEs, i.e. 53 in interzone and 35 in phalange, all CEs being located in the aforementioned *DACT2-SMOC2* region. Further, by then performing the analysis of H3K27ac, we showed that 45 (out of a total of 59) DacRs were significantly enriched in interzone. These differences in numbers of CEs as well as overall acetylation level partially explain the dynamics of 3D organization within the *DACT2*-*SMOC2* region, which presents higher frequency of *cis*-proximities in interzone than in adjacent phalange.

By integration of the 3D chromatin structure and H3 signatures data, we selected seven candidate enhancers (CE1 to CE7) located close to *DACT2*. Multiple studies have shown that the enhancers’ activities are associated with their states^55–57^. Thus, we used ChromHMM to characterize the states around these selected CEs, and annotated them according to the Roadmap Epigenomics Program^58^. We discovered interesting correlations, in particular the switching from strongly active to poised enhancer, and the interzone specific *cis*-proximity within the *DACT2* locus. Specifically, the chromatin state changes within the entire CE3, whereas CE1 and CE5 present a switch over a section of the enhancer region. Virtual 4C data revealed that CE7 presents *cis*-proximity with both *DACT2* and *SMOC2*, suggesting that this enhancer supports expression of more than one target gene.

Previous studies have shown that enhancer elements without homologous sequence in zebrafish can act as active enhancers in zebrafish^59–61^. This is supported by our findings, however, not all of the CEs were detectably active in the zebrafish assay, suggesting that some of the CEs can be chicken-specific. However, we were able to show that CEs1-3 and CE6 present activity during overall zebrafish development, which correlates with the known expression domain of *DACT2* in zebrafish larvae. Further studies in developing chicken embryos themselves will be required for detailed characterization of all *DACT2* and *SMOC2* enhancers in the interzone region.

## Experimental procedures

### Tissue collection

The experiments with mouse and chicken early embryos were performed in accordance with the relevant guidelines as applied and approved by the Ethical Committee at the Medical University in Lublin, where this work was performed, and also comply with the European regulations (directive 2010/63/EU).

Fertilized eggs (White Leghorn) were incubated at 38.5°C and 70% humidity for 7.5 days in a Grumbach BSS420 CTD7 incubator. The Hamilton Hamburger (HH) developmental stage was evaluated under Zeiss Stereo Discovery V8 microscope equipped with 0.63x Plan Apo S Objective Lens. Selected embryos at HH32 were sacrificed for tissue microdissection. The joint interzones and phalanges were collected from 2^nd^ and 3^rd^ hindlimb digit using a Dumont No.5 forceps (tip dimensions: 0.005 x 0.025 mm).

CD1 mice were group-housed in conventional cages conforming to local and (inter)national Animal Welfare Guidelines. Pregnant mice were sacrificed after 14 or 14.5 days by cervical dislocation. Subsequently, E14 or E14.5 mouse embryos were collected for RNA *in situ* hybridization.

### RT-qPCR

The Syngen Tissue RNA Kit was used for total RNA extraction, followed by treatment with DNase, using QIAGEN RNase-Free DNase Set. The RNA in biological triplicates for interzone and phalange was used for cDNA synthesis with Invitrogen™ SuperScript™ IV Reverse Transcriptase and Oligo(dT) primer, following the manufacturer’s suggestions. The qPCR was performed with LightCycler^®^ 480 Instrument II using PowerUp™ SYBR™ Green Master Mix II. Sequences of primers are listed in **Table S8**. Gene expression was normalized to expression of GAPDH. The T-test was used to analyze the expression of candidate genes.

### ChIP-Seq data analysis, candidate enhancer (CEs) identification and characterization of chromatin states

The ChIPseq data for H3K27ac and H4K4me1 was used from Nowosad et. al. (2022). The quality of raw fastq files was validated using FastQC (https://www.bioinformatics.babraham.ac.uk/projects/fastqc/). Next, reads were mapped to the *Gallus gallus* 5.0 reference genome using Bowtie 2 with default parameters^62^ followed by removal of PCR-duplicates using Picard (http://broadinstitute.github.io/picard/). The peaks were called using MACS2 with input as a control^63^. Default parameters and significance level threshold FDR <0.05 were used for MACS2 peak calling.

The differentially acetylated regions (DacRs) were identified based on the H3K27ac ChIP-Seq data using DiffBind tool (https://bioconductor.org/packages/DiffBind/) with default setting, except dba.count(summits = 1000). The log2FoldChange for identified DacRs were calculated using DiffBind binding affinity analysis following the default settings. Subsequently, the DacRs were filtered for FDR <0.05 and coordinates encompassing the *SMOC2-DACT2* region (chr3:40,15-43,6 Mb). The enrichment tracks for H3K4me1 and H3K27ac ChIP-Seq data were generated using deepTools2^64^. Briefly, bamCoverage with kilobase per million mapped reads (RPKM) normalization was used. The enrichment tracks were visualized using in-house R script.

Candidate enhancers (CEs) were identified using an in-house made script. Specifically, the consensus peaks called by MACS2 were defined by merging replicates using BEDTools. Next, the H3K27ac regions were intersected from H3K4me1 peaks using BEDTools intersect^65^, and promoter regions (−2.5/+2.5 kb from TSS) were removed from the dataset using GenomicRanges^66^.

### Targeted chromatin capture (T2C)

T2C protocol was further adapted from Boltsis *et al*. (2021)^39^. For each sample, 100 interzones or transient cartilages were dissected, respectively pooled, to obtain 10^6^ cells per tissue. The preparation of single cell suspension and chromatin crosslinking was performed as described in the ChIP protocol. Nuclei were extracted during 20 min by incubating the cells in ice-cold lysis buffer (10 mM Tris-HCl pH 8.0, 10 mM NaCl, 0.5% IGEPAL^®^ CA-630 and complete protease inhibitors) at 4°C. Isolated nuclei were washed twice by resuspension in 500 μl of PBS, followed by slow-spin centrifugation at 340 g, at 4 °C. Next, the nuclei in the pellet were resuspended in freshly prepared 1.2x restriction buffer, followed by addition of 10% SDS to a final concentration of 1.6% SDS and 1 hour of incubation at 37oC, while shaking at 900 rpm. SDS was quenched by addition of 20% Triton X-100 (final concentration: 1%) and incubation for 1 hour at 37°C while shaking at 900 rpm. The chromatin was digested with *ApoI* (New England Biolabs) (40 U/sample) for 16 hours at 37°C, while shaking at 900 rpm. The digested chromatin was ligated with T4-DNA Ligase (10 U/sample) at 16°C for 16 hours. The next day, chromatin was incubated with 3 μl of Proteinase K (10 mg/ml) for 1 hour at 65°C, and 3 μl of RNAse-A (10 mg/ml) for 45 min at 37°C, followed by DNA purification using Phenol:Chloroform according to the manufacturer’s instruction. The re-ligated DNA was digested with *DpnII* (50 U/sample) at 37°C for 16 hours, while shaking at 500 rpm, and purified by Phenol:Chloroform prior to T2C library preparation.

### T2C library preparation

The protocol was again further adapted from Kolovos and co-workers^38^ with previously applied modifications^39,40^. For the joint interzone and adjacent phalange sample, a T2C library was prepared using 350 and 175 ng of linearized chromatin, respectively. The samples were re-buffered to 10 mM Tris-HCl, pH 8 by a standard AMPure XP (Agencourt) bead clean-up procedure. The chromatin was sheared to 250-400 bp-sized fragments by a S220 Covaris (Covaris Inc.). The concentration was determined by Quant-it high sensitivity (ThermoFisher Scientific). For each sample, 100 ng of sheared chromatin was end-repaired and A-tailed using the Kapa hyper prep kit (Roche) according to the manufacturer’s instructions. SeqCap library adaptors were ligated followed by AMPure bead clean-up. The pre-capture library was amplified by PCR using KAPA HiFi hotstart readymix for 9 cycles. The amplified pre-capture library was purified by bead clean-up and quantified by Bioanalyzer DNA1000 assay (Agilent) according to the manufacturer’s instructions.

A *DACT2 – SMOC2* locus *Gallus gallus* 5.0-based design was ordered at NimbleGen (Roche) with baits located between 40,154,526 and 43,603,576 of Chr 3. A pooled hybridization mixture was prepared with 1 μg pre-capture library of each sample, 1 mM HE-index-oligo, 1 mM HE universal oligo, COT human DNA, AMPure XP reagent and added to 4.5 μl of pre-ordered baits and subsequently hybridized for 16 hours at 47°C. Post-hybridization, the samples were washed according to the instructions in the Nimblegen SeqcapEZ Hypercap workflow (Roche), the chromatin captured using capture beads. The captured library was amplified by PCR using Kapa HiFi mix and purified by AMPure XP beads. The captured library was quantified by Nanodrop spectrophotometer and the quality was assessed using a Bioanalyzer DNA1000 assay. Finally, the captured T2C libraries were denatured and sequenced on an Illumina HiSeq2500 sequencer with a custom recipe of 6 dark cycles, followed by paired end 101 sequencing with single index using the rapid v2 chemistry according to manufacturer’s instructions (Illumina) to a depth of approximately 30M clusters per sample.

### T2C data analysis

The analysis was performed using the pipeline described by Kolovos *et al*. (2018)^38^. Specifically, the quality of raw fastq files was evaluated using FASTQC. The reads were trimmed for adapters using AdapterTrimmer (https://github.com/erasmus-center-for-biomics/AdapterTrimmer) and mapped to the *Gallus gallus* 5.0 reference genome with the BWA aligner and the BWA-backtrack method. Alignments were subsequently annotated with the restriction fragments in which they were located. The proximity matrix was then constructed from the mapped primary alignments with their mapped primary mates. Further analyses and filtering based on the proximity matrix were performed in the R environment for statistical computing. Samples were normalized using array normalization^38^. The virtual 4C tracks were generated using in-house R script.

### Cloning and zebrafish transgenic enhancer assay and luciferase assay

Primers were designed to amplify CEs from chicken DNA (**Table S8**). Forward primers contain CACC flanking sequence complementary to the GTGG overhang sequence of the pENTR^™^/D-TOPO^®^ vector. The CEs were PCR amplified using touchdown PCR protocol with Phusion^™^ High-Fidelity DNA Polymerase. The PCR products were cloned into pENTR^™^/D-TOPO^®^ vector using pENTR^™^ Directional TOPO^®^ Cloning Kit, followed by recombination to Zebrafish Enhancer Detection (ZED) vector^43^. The ZED vector contains green fluorescent protein GFP under GATA minimal promoter and red fluorescent protein (DsRed2) under cardiac actin promoter.

The wild-type strain zebrafish larvae were handled according to the welfare regulations and standard protocols approved by the Ethical Committee at the Medical University in Lublin, which comply with the European regulations (directive 2010/63/EU). The CEs/ZED and empty ZED vector alone were injected using standard procedures under the Stemi 508 stereo microscope (ZEISS) equipped with a microinjection unit. Specifically, the single-cell stage eggs were used for injection of 1nl mixture containing 1:1 ratio of ZED (50 ng/μl) along with Tol2 transposase mRNA (40 ng/μl) to facilitate genomic integration^59^. For statistical significance, the 130 to 200 embryos were injected with each construct, and for each construct a full experiment was conducted at least twice to ensure reproducibility and high number of embryos injected. Eggs that were unfertilized or damaged during the injection process were removed after approximately 10 hours post-injection, as their presence could negatively affect the survival of remaining embryos. Larvae were then maintained in standard conditions (28.5°C) and the efficiency of the genomic integration was validated based on the DsRed2 expression pattern analyzed after 72 hours. The 1-phenyl 2-thiourea (PTU) at the standard concentration of 0.003% (200 μM) was added to the E3 medium to delay pigmentation and therefore more accurately visualize the fluorescent signals. The specimens without DsRed2 expression in muscles (positive control of construct integration) were excluded from further examination. Next, the larvae were screen for expression of GFP after 96h. If GFP expression pattern was present in at least 30% of specimens, it was considered as enhancer-specific. Larvae were screened for expression patterns under the SteREO Discovery.V8 fluorescence stereomicroscope (ZEISS, Germany) and images were captured using the ZEN2 software (ZEISS, Germany).

## Data availability

The T2C data is available under the gene expression omnibus (GEO) accession number GSE210108.

## Supplementary data

The supplementary data is available at https://github.com/karolnowosad/Chromatin-architecture-and-cis-regulatory-landscape-of-the-DACT2-SMOC2-locus-in-the-developing-synov.git

## Acknowledgments

We would like to thank the members of the Center for Biomics-Genomics at the Erasmus Medical for technical support and the Experimental Medicine Center at the Medical University of Lublin for taking care the mice and zebrafish.

